# Landscape matters: Deriving a generalizable understanding of population connectivity using empirical data and graph theory

**DOI:** 10.1101/2025.09.05.674386

**Authors:** Divyashree Rana, Samuel Alan Cushman, Uma Ramakrishnan

## Abstract

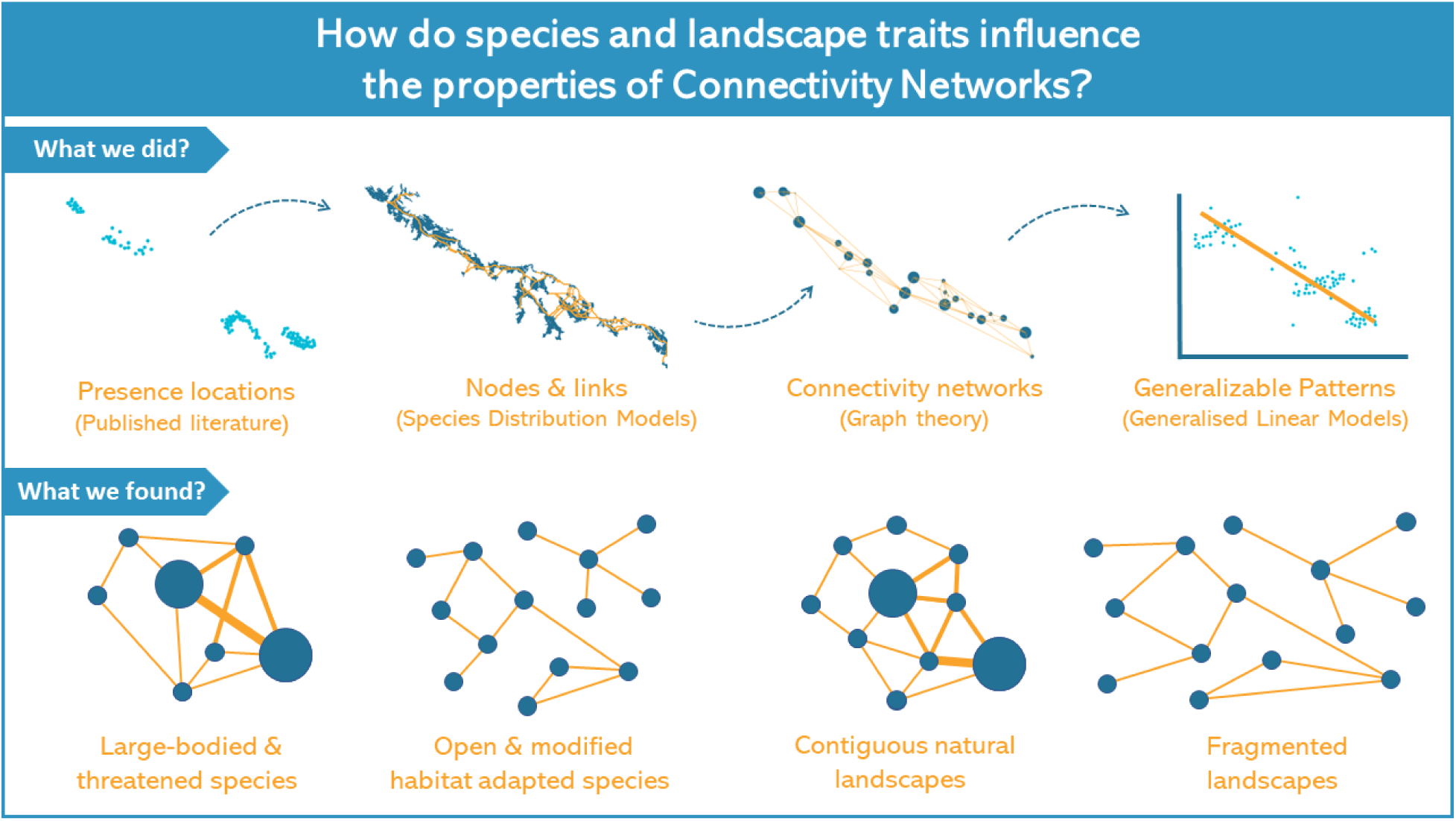

Human impacts on ecosystems have accelerated globally, driving a 10% decline in terrestrial biodiversity and a 70% decline in wildlife populations over the past five decades. These losses are closely linked to habitat modification and fragmentation, highlighting the urgent need for management strategies grounded in a clear understanding of how wildlife use landscapes and navigate human-altered areas. Connectivity between populations is critical for species persistence and is shaped by the interplay between landscape features and species movement. Most connectivity studies pursue application-focused goals, such as designing corridors or assessing land-use effects, often targeting single species within specific landscapes. While these approaches provide depth, they limit the development of general principles that apply across species and regions. Graph theory offers a powerful framework to distill complex connectivity patterns into comparable metrics, creating opportunities to identify such generalities.

In this study, we examined connectivity patterns for 11 Indian carnivore species distributed across heterogeneous landscapes. Using secondary data, we developed habitat networks from species distribution models that incorporated both habitat quality and matrix resistance. We then applied graph theory to generate networks based on connectivity between identified habitat nodes, enabling comparisons across species and landscapes. Our results show that while connectivity patterns differ markedly among species, broad trends emerge. Larger-bodied species like tigers, which in our study are often threatened species, can overcome the effects of fragmentation better than smaller bodied species, however their connectivity is dependent on the existence of high-quality patches. Fragmented and heterogenous landscapes were always associated with modular, less efficient networks irrespective of the species. Importantly, landscape characteristics had greater influence on network-level connectivity properties, while species traits more strongly determined node-level structural complexity of the network. By integrating network theory with multispecies analyses across diverse landscapes, our work moves beyond single-species case studies to identify general drivers of connectivity. In terms of conservation, our approach allows us to generate broad insights into drivers of population connectivity informing strategies for lesser-known species and guide more effective, landscape-scale management in an era of rapid environmental change.

## 1. Introduction

Habitat loss and fragmentation are driving population declines of many species globally (Tucker et al., 2018). The ability to navigate and utilise modified landscapes likely strongly influences the persistence of species in the future (Goodwin & Fahrig, 2002; Baguette et al., 2013). Hence, understanding species movement determining population connectivity is a fundamental topic for conservation researchers and practitioners alike (Crooks and Sanjayan, 2006). Since the 1990s, research in connectivity conservation has seen a sharp increase, however, most of these studies are regional in scope (Correa Ayram et al., 2016). Much of the wildlife connectivity research is case-specific focussing on single or few species within a landscape. Despite growing evidence of similar effects of barriers like roads on the connectivity of multiple species (Fu et al., 2010; Thatte et al., 2019), there is a lack of research trying to assess the existence of any generalizable patterns in connectivity research. There remains little synthesis of whether consistent, generalizable rules link species and landscape traits to connectivity (Liczner et al., 2024). Identifying such rules could transform scattered case-specific insights into a broader theoretical framework, while also guiding conservation in data-deficient systems (Schaffer-Smith et al., 2016).

Connectivity across populations is a crucial determinant of species persistence in any landscape (Wang et al., 2021) and depends on the interaction between landscape features and species movement (Taylor et al., 1993; Baguette et al., 2013). Population connectivity is driven by i) species traits that determine their habitat suitability and dispersal abilities, and ii) properties of the landscape matrix facilitating or hindering species movement (Goodwin & Fahrig, 2002). Graph theory offers a powerful framework for translating complex ecological landscapes into simple, quantitative representations of connectivity (Urban et al., 2009). By abstracting habitat patches and their linkages into nodes and edges, graph-based metrics capture key properties of networks, such as efficiency, clustering, and centrality, that are directly relevant to movement and gene flow (Keeley et al., 2021; Hashemi & Darabi, 2022). Importantly, these metrics are comparable across species and landscapes, making graph theory uniquely suited for identifying general principles rather than site-specific outcomes. While previous studies have used graph-based approaches in single-species or local contexts, their potential to reveal broader, trait-based patterns of connectivity across taxa and landscapes remains underexplored.

Can consistent species or landscape properties predict connectivity outcomes across contexts? Addressing this question is crucial, as it could help uncover the underlying processes that shape connectivity across ecological contexts while also offering a framework to guide conservation strategies for data-deficient or unstudied species. There has been a theoretical advancement in understanding the role of landscape structure (Goodwin & Fahrig, 2002), matrix characterization (Bender and Fahrig, 2005; Koenig & Bender, 2018), followed by fewer methodological comparisons in measuring connectivity (Wade et al., 2015; Poli et al., 2022; Lumia et al., 2024; Schippers et al., 2025). Most of these studies are either based on simulation (Koenig & Bender, 2018; Ash et al., 2020) or experimental data (Poli et al., 2022) or are built on small empirical datasets (Borthagaray et al., 2012). The narrow context of case-specific study provides detailed insights but limits identification of overarching principles uncovering consistent landscape and species correlates of connectivity patterns.

In this study, we assess trends in the connectivity of carnivores across India as a function of species and landscape traits using graph theory on a large empirical dataset. Carnivores present a good test case as they are naturally low-density, wide-ranging, sensitive to fragmentation and hence often of conservation priority (Dinerstein et al., 2007; Borthagaray et al., 2012). By testing these predictions on a diverse carnivore assemblage across Indian landscapes, we explore whether generalizable patterns of connectivity can be identified at macroecological scales. This involved testing multiple predictions - i) larger species will have more connected, large-scale networks across landscapes, ii) threatened or rare species will exhibit more fragmented and fragile networks, with limited connectivity and higher modularity, iii) species adapted to open, modified landscapes will have more resilient and redundant networks, iv) more continuous and cohesive landscapes will support simpler, highly connected networks with fewer isolated nodes or modules, v) highly fragmented landscapes will result in modular networks with many isolated patches or subnetworks, and vi) landscapes with high habitat heterogeneity will promote structurally complex and modular networks. Understanding such patterns would not only build on our limited understanding of generalizable principles governing carnivore connectivity but could also help in conservation decision-making for data-deficient species.

## 2. Methods

We generated connectivity networks to understand the role of species and landscape traits in determining network properties using Generalized Linear Models (GLMs). Incorporating graph theory, our methodological workflow consisted of four major steps: a.) identification of suitable habitat patches to be classified as nodes, b.) estimation of species-specific resistance surface, c.) identification of weighted links connecting nodes, and d.) generation of species and landscape-specific connectivity networks. The first two products were achieved from the habitat suitability predictions generated from Species Distribution Models (SDMs) built for each species across India. The resistance surface was used to identify least-cost paths or links between nodes. Finally, the nodes and links together were used to generate connectivity networks for 11 species of carnivores across India. These steps have been detailed below.

### 2.1. Species presence records and traits

To generate comparable networks across the country, 11 species of carnivores were selected - *Panthera tigris* (tiger, TG), *Melursus ursinus* (sloth bear, SB), *Panthera pardus* (leopard, LP), *Hyaena hyaena* (striped hyena, SH), *Canis lupus* (Indian wolf, IW), *Canis alpinus* (dhole, DH), *Canis aureus* (golden jackal, GJ), *Prionailurus viverrinus* (fishing cat, FC), *Felis chaus* (jungle cat, JC), *Prionailurus bengalensis* (leopard cat, LC), and *Prionailurus rubiginosus* (rusty-spotted cat, RSC). The selection of carnivores was made to have species with wide overlapping distributions, and varying body size, dispersal limits, and habitat preferences. Presence locations for selected carnivores were extracted from published literature using the studies collated in Srivathsa et al. (2020) and Jhala et al. (2020). Only the sources that provided exact coordinates or maps that could be georeferenced were used to extract occurrence records, following Rana et al. (2024). We then spatially thinned these records by applying a circular buffer of the size of each species’ home range (Table S1) using the *spThin* package in R (Aiello-Lammens et al., 2015) to minimise georeferencing error as well as bias induced due to uneven sampling effort (Kramer-Schadt et al., 2013).

### 2.2. Building species distribution models

The predictor variables were retrieved and rescaled to 1 sq km from multiple datasets in Google Earth Engine to include climatic, biological, geomorphological, and anthropogenic variables affecting species distributions (details in Table S2). Ecologically relevant layers which were not highly correlated (r < |0.7|) were retained. These layers were used to independently build Species Distribution Models (SDMs) for each species. Considering the sensitivity of SDMs to predicting outside the training environmental niche space (Yates et al., 2018; Rana et al., 2024) and the heavy computational demands of the analysis, the data were partitioned to build independent species distribution models and connectivity networks within smaller grids (5×5 degrees) for each species. The grids with fewer than 15 spatially thinned species records were dropped from further analysis. Following this, within each grid, pseudo-absences ten times the occurrence records were generated for training the SDMs. Given the wide distribution of the selected species and habitat heterogeneity in India, the smaller spatial scale of the analysis ensured more accurate and localized habitat predictions while minimizing predicting outside the training extent.

There are several algorithms available and recommended to model the distribution of species using presence-only data. However, ensembles of tuned individual algorithms have been shown to provide high predictive power (Valavi et al., 2022). Hence, for all species, independent SDMs with a set of uncorrelated predictor variables (r<|0.7|) within each grid were built using six algorithms. These included machine learning algorithms - Maximum Entropy Modelling (MaxEnt) and Support Vector Machines (SVM), tree-based algorithms - Boosted Regression Tree (BRT), Random Forest (RF), and regression algorithm - Generalized Additive Models (GAM) and Multivariate Adaptive Regression Splines (MARS). Model performance is highly sensitive to the parameters of each algorithm, advocating for individual model tuning instead of relying on default settings (Hallgren et al., 2019; Valavi et al., 2022; Soley-Guardia et al., 2024). Hence, model parameters were tuned for each algorithm using *ENMeval* and *caret* packages in R (Kuhn & Johnson, 2013; Muscarella et al., 2014). Predictive models were built with optimised parameters and validated through a random ten-fold cross-validation approach for each algorithm using the *sdm* package in R (Naimi & Araújo, 2016). Lastly, AUC-weighted ensemble predictions were generated using all optimized models. Hence, for each species, an ensemble habitat suitability prediction was created within each grid. Further details of the SDM modelling approach can be found in Samad et al. (2024).

### 2.3. Generating connectivity networks

As explained earlier, habitat suitability predictions from the SDMs were used to identify suitable habitat patches (classified as nodes) as well as to generate resistance surfaces within each grid for all species. Using the identified nodes and resistance offered by the interspersed matrix, least-cost paths (LCPs, edges) were identified between each node pair. Once habitat patches and LCPs were identified, these species-specific networks were translated into simpler spatial graphs (as represented in Figure 1) with nodes and links with their associated strengths and attributes, henceforth called “connectivity networks”. Details of the workflow can be found below.

**Figure 1.**
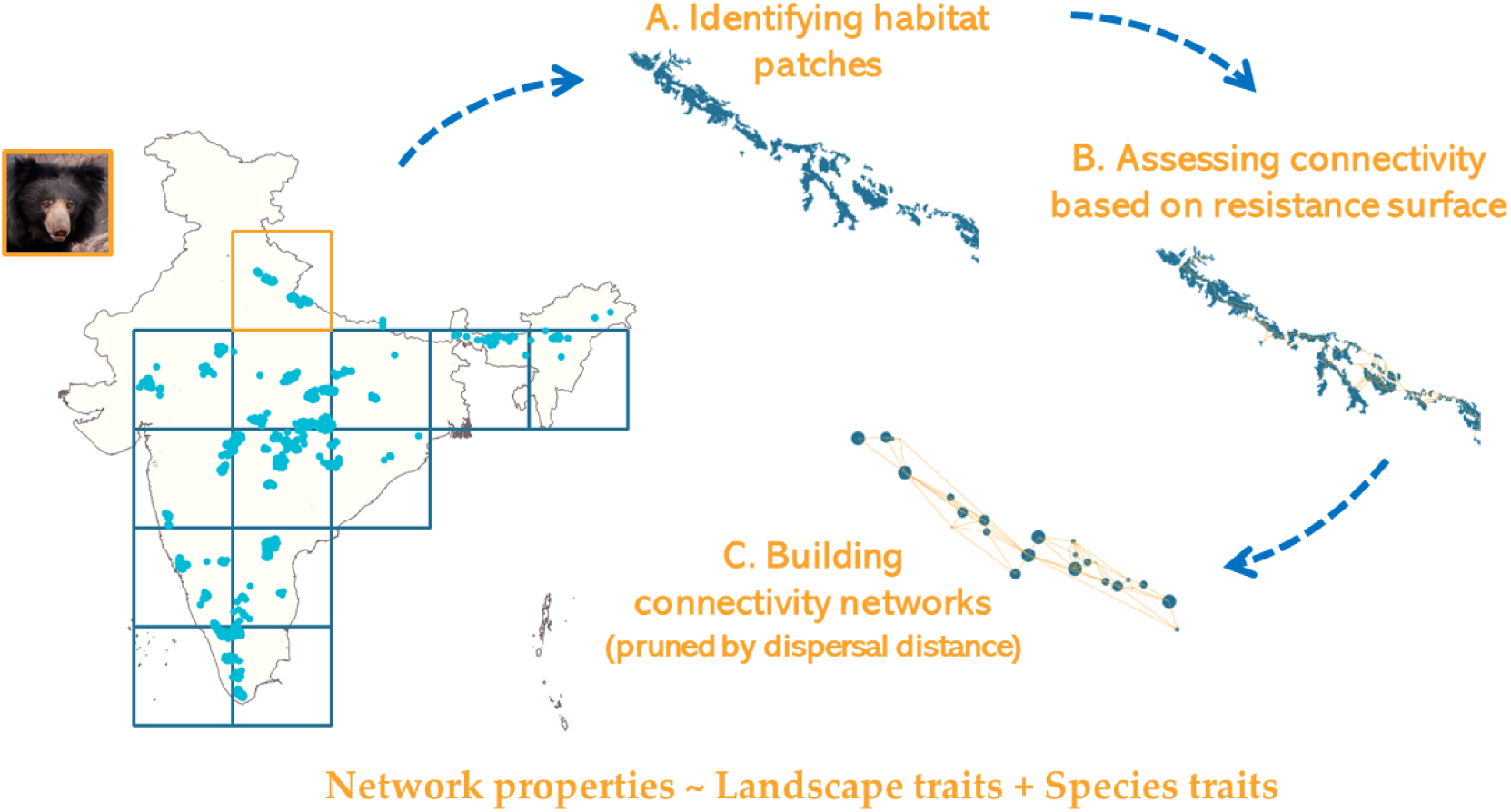
Schematic for methodological workflow for generating connectivity networks with an example of sloth bear (*Melursus ursinus*) in Terai landscape. A. Identifying habitat patches limited by home range size using SDM predictions, B. Generating habitat networks with LCPs based on resistance matrix, and C. Generating spatial graphs from habitat networks.

Firstly, to identify nodes, a binary habitat/non-habitat raster was generated from habitat suitability predictions based on the threshold maximizing the sum of sensitivity and specificity of the model (Max SSS) (Liu et al., 2013). Using this layer, nodes were identified based on species-specific patch size threshold (larger than average home range, Soria et al., 2021). Next, a resistance surface using a negative exponential transformation of the habitat suitability values ranging from 1 to 100 (c8 transformation, Keeley et al., 2016). Missing values in a raster were replaced with the highest resistance value. Using the resistance surface, a cost-based linkset was created between the identified nodes to generate complete topography spatial graphs using the *graph4lg* package in R (Savary et al., 2020). These complete graphs were pruned based on the species-specific thresholds to remove any edges with lengths more than the dispersal limits of the species using the *igraph* package in R (Csardi & Nepusz, 2006). This generates pruned connectivity networks incorporating species ecology for realistic downstream inferences. Average species dispersal limits were incorporated based on data available in the COMBINE database (Soria et al., 2021), however, as dispersal limits could vary across individuals, sensitivity analysis with a 25% increase and decrease of the threshold was also conducted. These pruned graphs were used to calculate network properties to understand the generalizability of connectivity patterns across multiple species and landscape traits.

### 2.4. Extracting species traits, landscape traits and network properties

Species traits relevant to understanding network connectivity were collated across selected species from published literature (details in Table S2). Species-specific home range was used to filter out smaller nodes whereas dispersal distance was used to filter out longer links. Remaining species traits were used to understand the network properties. Similarly, landscape traits relevant to understanding resistance to movement were calculated for each 5×5 degree grid based on the landcover classification (ESA WorldCover 10m v100, Zanaga et al., 2021) using *landscapemetrics* package in R (Hesselbarth et al., 2019). These landscape traits were calculated independent of species, and for each grid. These species and landscape traits (details in Table 1) were used as predictors to understand their role in explaining various network properties. Highly correlated traits were removed to avoid multicollinearity in the dataset.

**Table 1:** Details of species and landscape traits to understand the properties of connectivity networks.

| Trait name | Trait type | Explanation | Source |
| --- | --- | --- | --- |
| Home Range | Species | Size of the area within which the everyday activities of individuals or groups of individuals are typically restricted in km<sup>2</sup> | Soria et al., 2021 |
| Dispersal distance | Species | The distance an animal travels from its place of birth to the place where it reproduces in kilometers |  |
| Adult body mass | Species | Body mass of an adult individual in grams |  |
| Population density | Species | Number of individuals of the species per squared kilometer |  |
| Open habitat preference | Species | Preference to open habitats like grassland, wetlands, agricultural fields on the scale of one (closed habitat specialist) to five (open habitat specialist) | IUCN Redlist Assessment |
| Anthropogenic adaptability | Species | Adaptability to anthropogenic landscapes on the scale of one (strictly avoids modified landscapes) to five (found in modified landscapes) | IUCN Redlist Assessment |
| IUCN status | Species | Threatened status coded on the scale of 1 (least concern) to 5 (endangered) | https://www.iucnr edlist.org |
| Number of patches | Landscape | Number of patches; does not account for landscape composition or configuration | *lsm_l_np* |
| Largest patch index | Landscape | Size of the largest contiguous land cover patch; | *lsm_l_lpi* |
| Edge density | Landscape | All edges in the landscape in relation to the landscape area | *lsm_l_ed* |
| Proportion of natural habitat | Landscape | Percentage of natural land cover within the grid | *lsm_c_pland* |
| Largest patch index | Landscape | Percentage of the landscape covered by the largest patch of natural land cover | *lsm_c_lpi* |
| Patch density | Landscape | Patch density in the landscape increases as the landscape gets patchier | *lsm_l_pd* |

On the other hand, the network properties were used as the response variables determining various network properties. These were extracted for each connectivity network within a grid for all species using the *igraph* package in R (Csardi & Nepusz, 2006). These properties were segregated based on two classification criteria: a. their spatial scale (network and node) and b. their function (structural complexity, connectivity, and centrality), as detailed in Table 2. Firstly, network properties were classified based on the spatial scale of analysis - properties calculated for the entire network as “network” and the properties calculated for each node and averaged for the entire network as “node”. Similarly, the properties were reclassified based on their functional role in explaining the features of a network. Properties that describe the basic architecture of the connectivity networks, like the number of nodes and edge density, were categorized under the function “structural complexity”. On the other hand, properties like network efficiency and mean flux capture how easily individuals/information can move through the network, and hence were categorized as determining “connectivity”. Lastly, the properties categorized as determining “centrality” assess the relative importance of nodes in a network, with higher values indicating relative higher importance of a few nodes in a network and lower values hinting at more redundant networks.

**Table 2:**
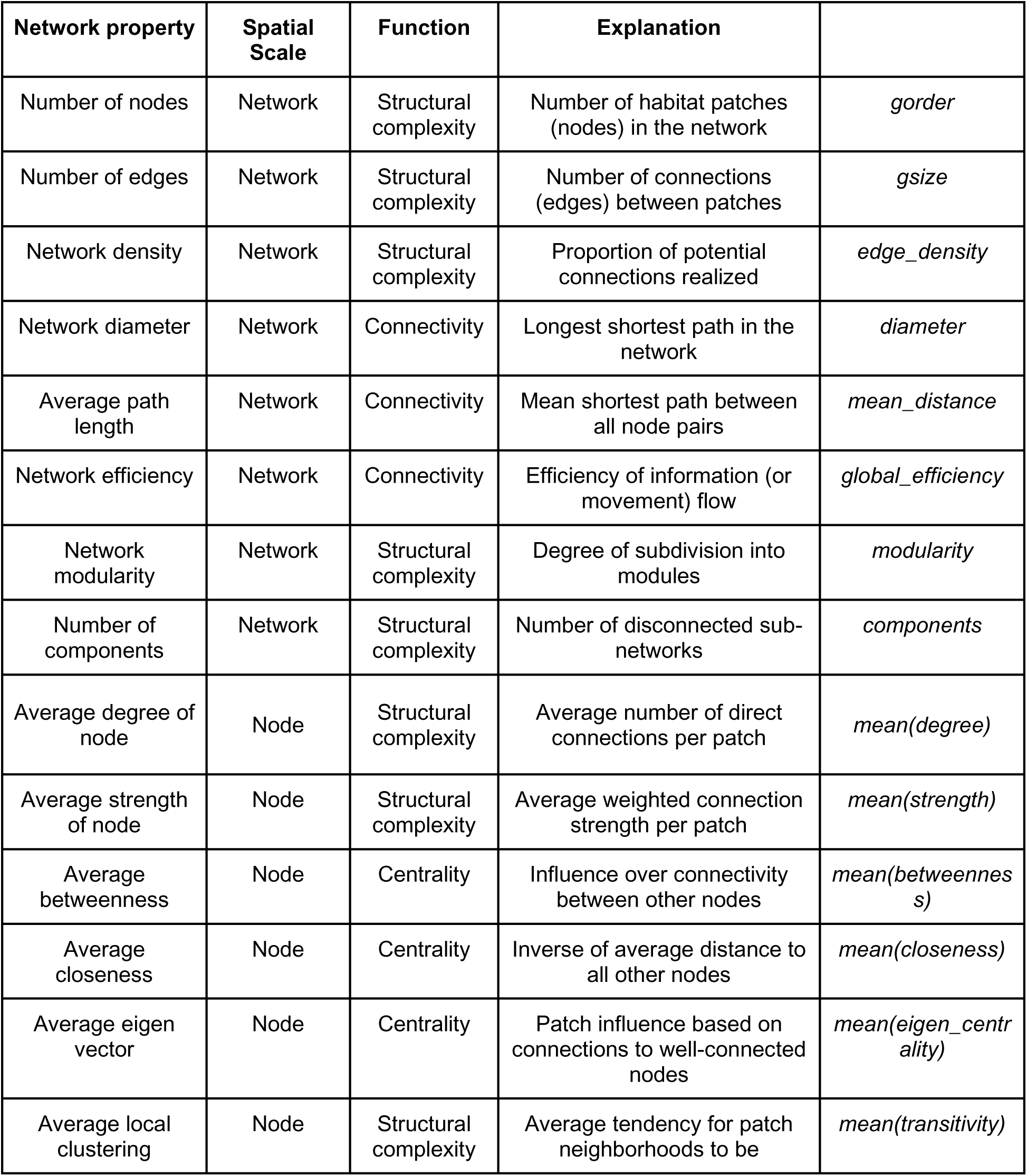
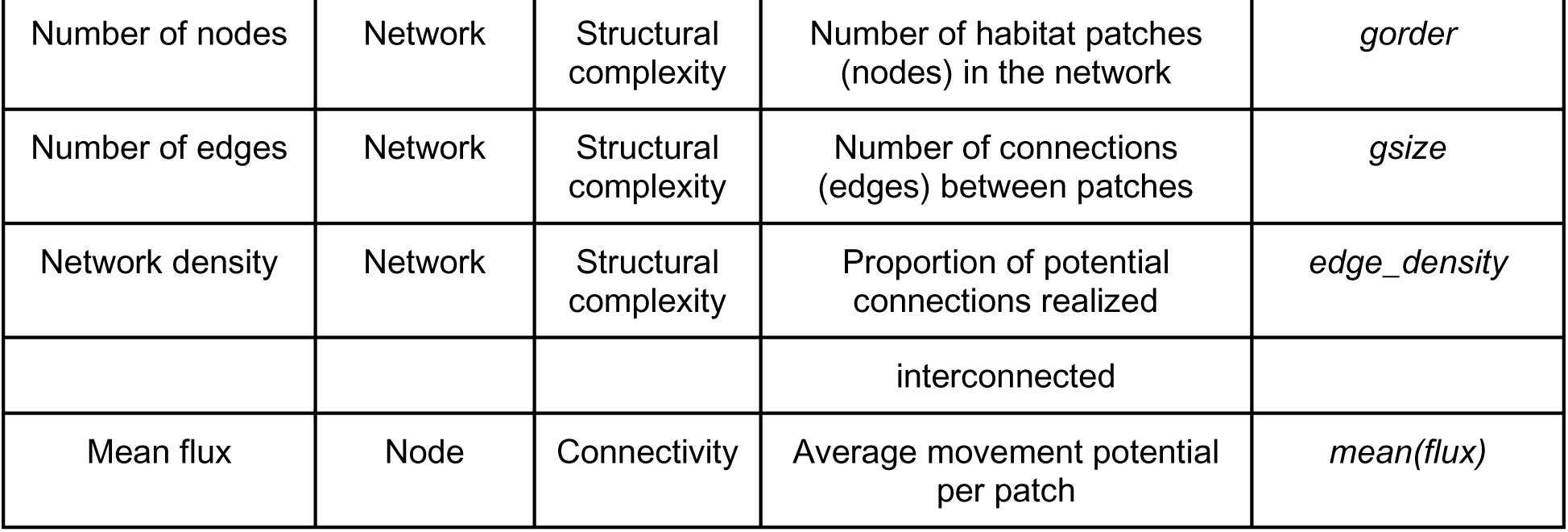
Details of network properties and their classification to understand the role of species and landscape traits in explaining them.

### 2.5. Understanding the role of traits in explaining network properties

GLMs were used to assess the relationship between network properties (response variables) and species and landscape traits (predictor variables). Models were implemented using the *glm* function in base R (R Core Team, 2023) and the *betareg* package (Zeileis et al., 2016). All predictor variables (species and landscape traits) were standardized (mean = 0, SD = 1), and response variables (network properties) were log transformed with a small offset to account for zeros. The choice of distribution family was guided by the characteristics of the response variable: if log-transformed responses approximated normality based on the Shapiro–Wilk test, a Gaussian family with identity link was used; if the responses were non-negative integers, a Poisson family was fitted; and if the responses were continuous, positive, and skewed, both Gamma and Beta regression models were fitted and the family with the lowest Akaike Information Criterion (AIC) was selected. Final models were evaluated for parameter significance (p < 0.05) and fit statistics (pseudo-R²) using the *performance* package in R (Lüdecke et al., 2021).

To examine both independent and combined effects of predictors, univariate models were fitted for each trait–property combination, and additive models were fitted with all predictors for each network property. Further, network properties were clubbed into two classification schemes, based on spatial scale and functionality as explained above, and the relative importance of each variable across multiple network properties was averaged to understand generalizable trends. The relative importance of predictor variables in additive models was estimated using a permutation-based approach, inspired by the *relaimpo* package (Grömping, 2007). For each network property, models were fitted across many random predictor orderings (n = 1000). The increase in model fit (pseudo-R²) attributable to each predictor was sequentially recorded within each ordering and averaged across all permutations to obtain the marginal contribution of each predictor, normalized to sum to one. This provided a robust estimate of relative predictor importance, generalized to the GLM families used in this study. These models allowed us to test our predictions to enrich our understanding of generalized patterns of connectivity networks with respect to species and landscapes.

## Results

### 3.1. Species and landscape correlates of network properties

To examine how species and landscape traits influenced connectivity, we fitted univariate GLMs between each trait and network property. Expected relationships based on the ecological predictions were compared against observed correlations. Results are summarized in a heatmap (Figure 2), which highlights the direction and strength of significant correlations, and highlights mismatches between prediction and observation with bold outlines for specific trait–property pairs. Below, we describe the major trends emerging from species traits and landscape traits separately.

**Figure 2.**
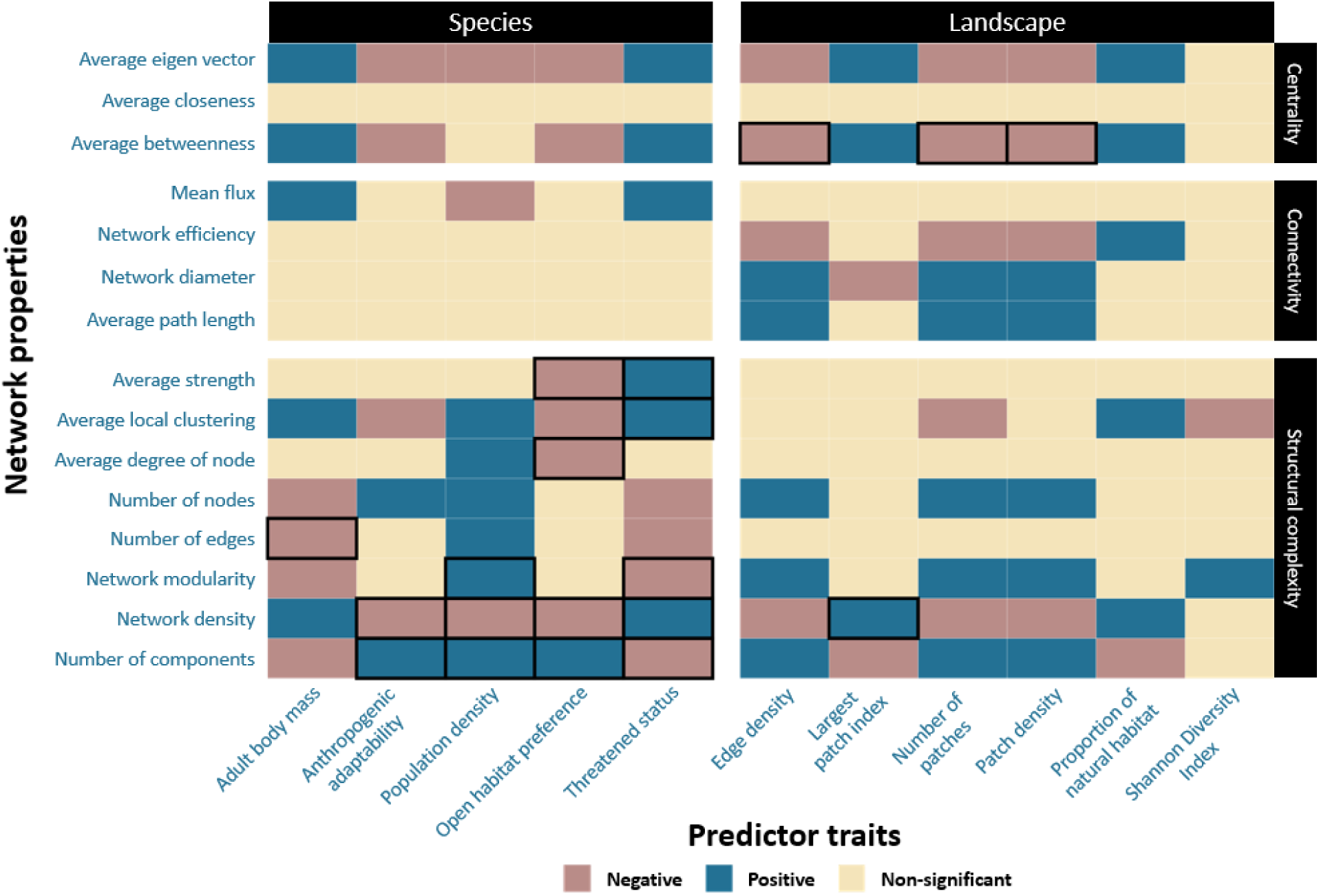
Plot showing the results of the GLM of network properties (y-axis) with species and landscape traits (x-axis), with cells coloured for significant relations (p<0.05) in positive or negative direction. Cells with black outline mark the significant trends which were opposite to our predictions outlined in Table S3.

Body mass was positively correlated with denser and large-scale networks with low modularity. Although networks of larger-bodied species contained fewer nodes, they exhibited higher local clustering within connected neighbourhoods. Larger body mass was also associated with more centralized networks, characterized by higher flow betweenness and eigenvector centrality. Species rarity, inferred from decreasing population density and increasing IUCN threatened status, revealed consistent patterns, indicating that both traits captured similar trends. Rare and threatened species were correlated with dense networks with fewer nodes and edges. These networks displayed strong connections within neighbouring patches and were also more centralized, with unequal patch importance. In contrast, species preferring open habitats or adapted to modified landscapes exhibited patterns inverse to those of threatened and rare species. Open-habitat species had sparse networks with multiple components, and although they contained more nodes, they exhibited lower local clustering, flow betweenness, and eigenvector centrality. Global network properties determining connectivity within the network did not show significant relations with any of the species’ traits.

Landscapes with higher proportions of natural habitat and larger patch sizes were positively correlated with denser networks that had greater efficiency and shorter diameters. Such contiguous habitats also supported more centralized networks, reflected in higher betweenness and eigenvector centrality. In contrast, fragmented landscapes, characterized by a greater number of patches, higher patch density, and higher edge density, were consistently associated with sparse and modular networks comprising multiple components. With increasing fragmentation, network efficiency declined, while diameter and average path length increased. These fragmented landscapes also showed lower centrality, with reduced betweenness and eigenvector values. Finally, heterogeneous landscapes, captured by high Shannon diversity of land-cover classes, were correlated with high modularity and reduced connectedness between neighbouring patches. However, landscape heterogeneity did not exhibit strong or consistent correlations with most other network properties, indicating more variable relationships.

### 3.2. Linking predictions with observed patterns

To evaluate the consistency of empirical patterns with theoretical expectations, six predictions linking species and landscape traits to network properties were summarized (Table 3). For each prediction, the associated trait and the corresponding observed pattern were recorded, providing a structured assessment of how species- and landscape-level factors aligned with expected effects on network properties.

**Table 3:**
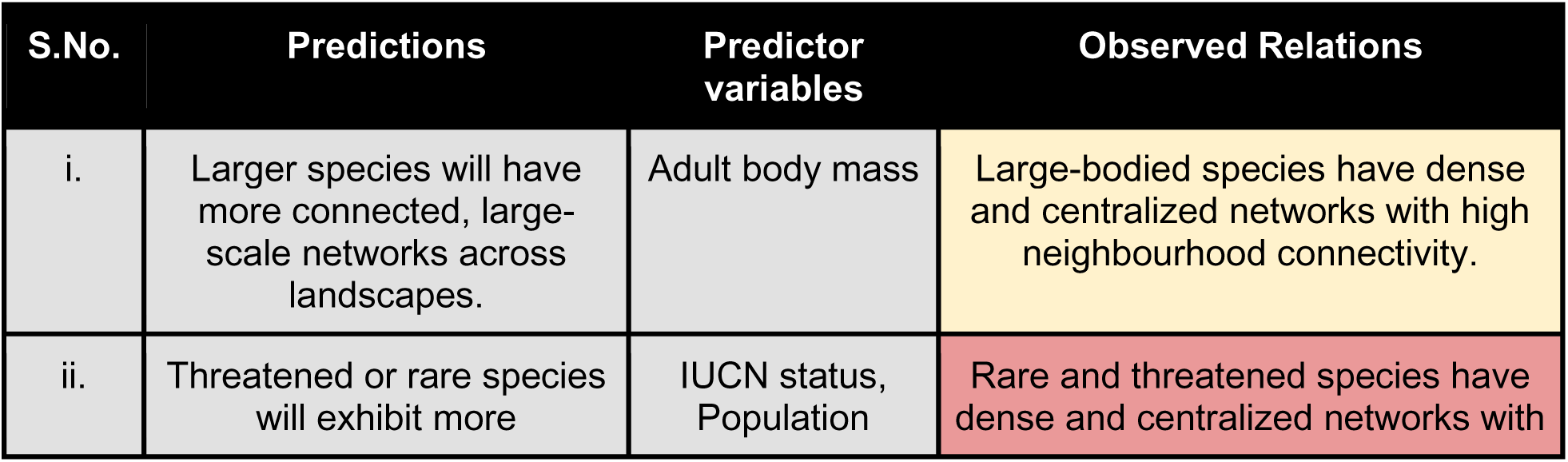
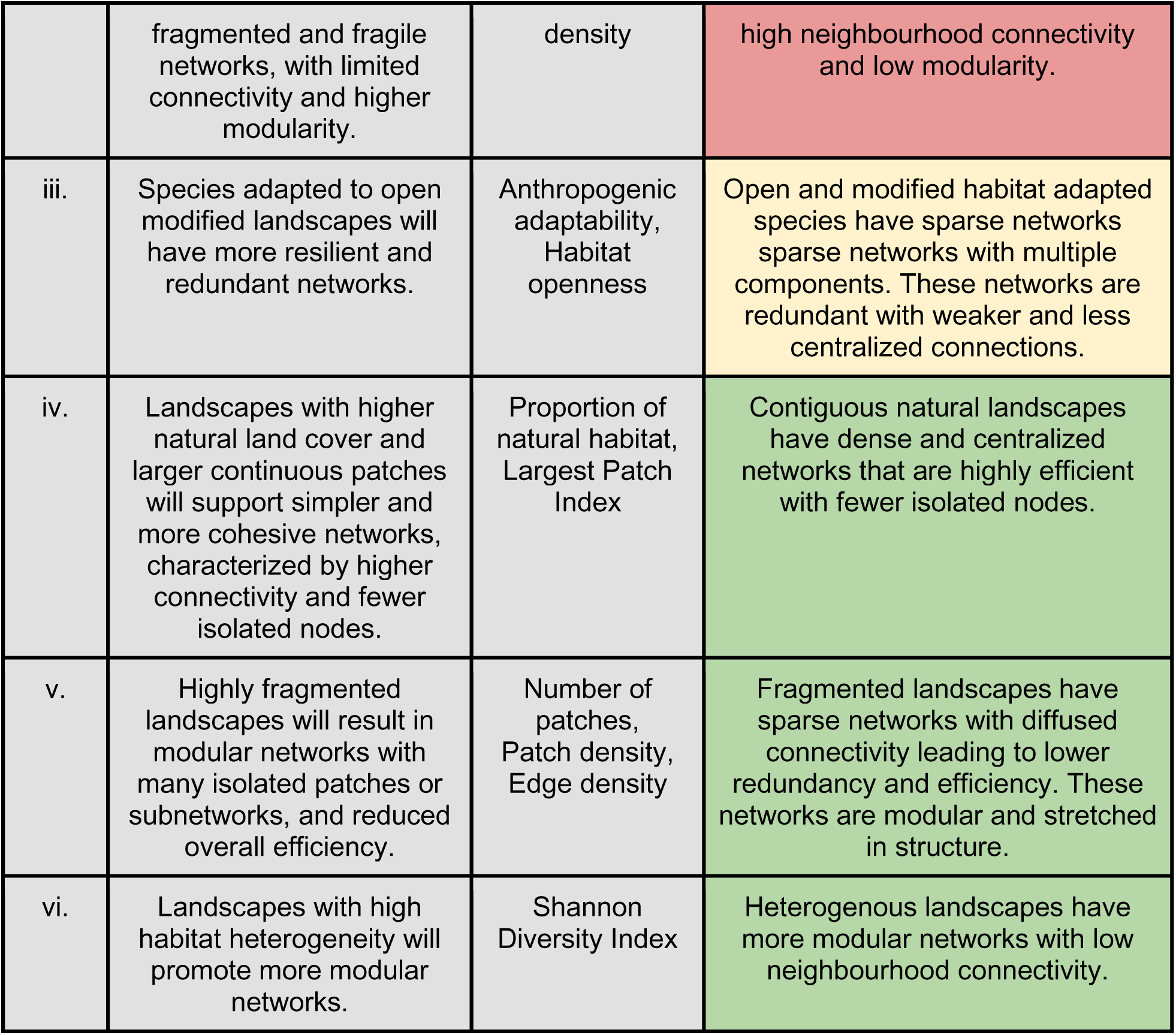
Predictions and observations for the relation between network properties with species and landscape traits and observed relations. The colour of the cell represents how well the observed relation matched the prediction - green (complete match), yellow (partial match), and red (mismatch).

Overall, most predictions were supported by observed patterns. Predictions related to body size and open-habitat preference aligned well with expectations, although large-bodied species exhibited dense but more centralized networks than anticipated, with uneven patch influence. In contrast, predictions for rare and threatened species showed major deviations, as they also formed dense networks instead of the expected modular and fragmented structures. Species traits generally showed non-significant associations with connectivity-specific metrics such as network efficiency (Figure 2), limiting a direct match to predictions. By comparison, landscape trait predictions corresponded closely with observations, indicating that mismatches were largely confined to species-level expectations (Figure 2).

### 3.3. Relative trait contributions to network properties

In addition to univariate relationships, we fitted additive GLMs to assess the relative contribution of species and landscape traits in explaining connectivity patterns. Response variables were grouped by their scale of estimation, network-level (global) and node-level (local), and the differences in trait contributions were compared (Figure 3). Node-level properties, indicated by negative relative differences, were primarily influenced by species traits (light bars), with open habitat preference emerging as the strongest predictor. In contrast, network-level properties were predominantly governed by landscape traits (dark bars). Among these, patch density, proportion of natural habitat, edge density, largest patch index, and number of patches contributed most strongly to shaping global connectivity. Shannon diversity index was an exception among landscape traits, contributing more substantially to node-level properties. These results highlight a scale-dependent influence of traits, where species attributes shape local connectivity while landscape context drives emergent network-level structure.

**Figure 3.**
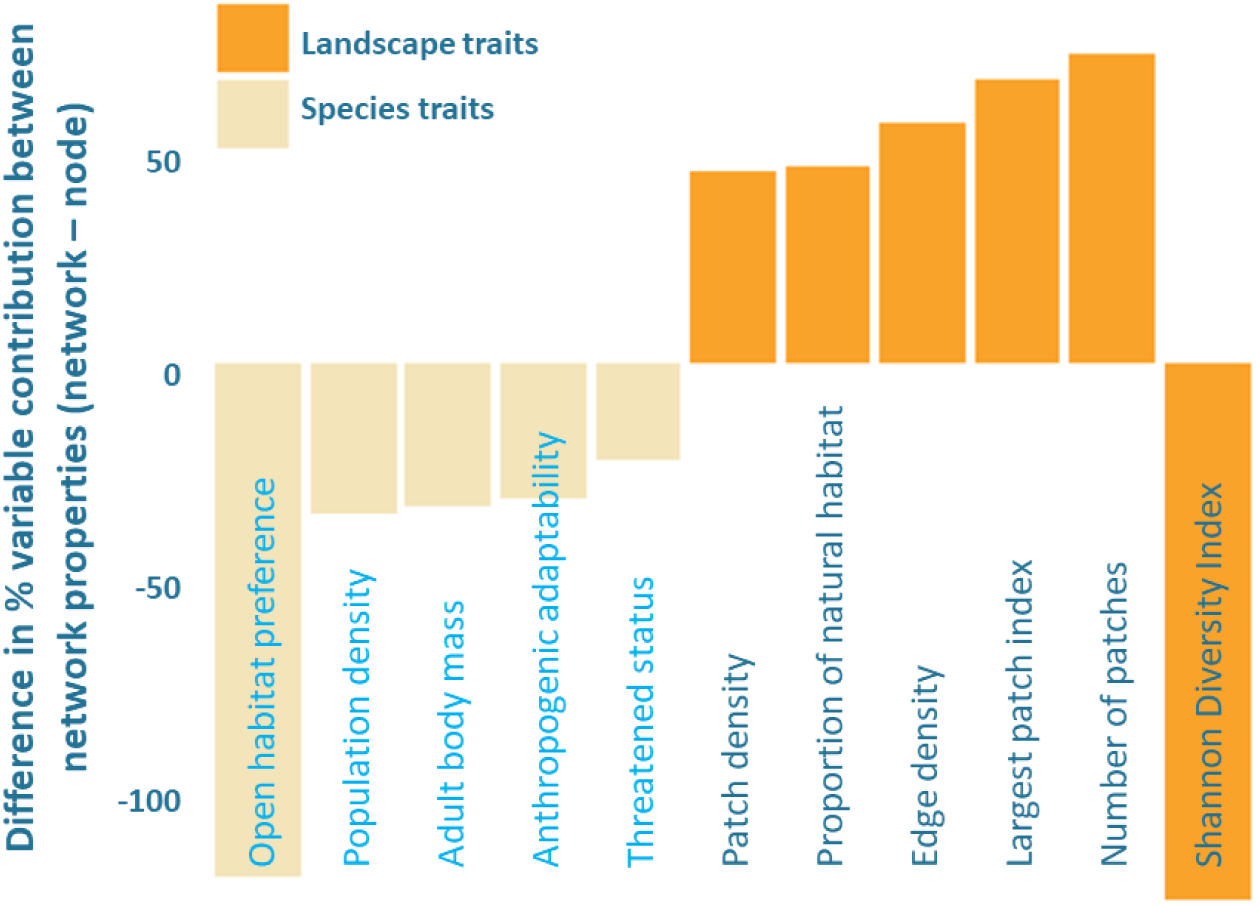
Difference in percentage variable contribution of species (light) and landscape (dark) traits in explaining network properties (network - node). Negative values indicate lower relative contribution in explaining network level properties as compared to node level properties.

In addition to categorizing network properties based on their scale of estimation, we further grouped them according to their functional role. Relative contributions of species and landscape traits to these functional categories revealed distinct patterns (Figure 4). Connectivity metrics (dark purple bars) were almost entirely explained by landscape traits, with negligible contributions from species traits. In contrast, species traits showed consistently higher contributions to properties determining centrality and structural complexity. Among species traits, open habitat preference, adult body mass, and threatened status explained a greater share of variation in centrality-related properties. Within landscape traits, patch density and Shannon diversity were dominant in shaping connectivity-related properties. Overall, the partitioning of contributions highlighted a functional divide between landscape and species traits across network properties.

**Figure 4.**
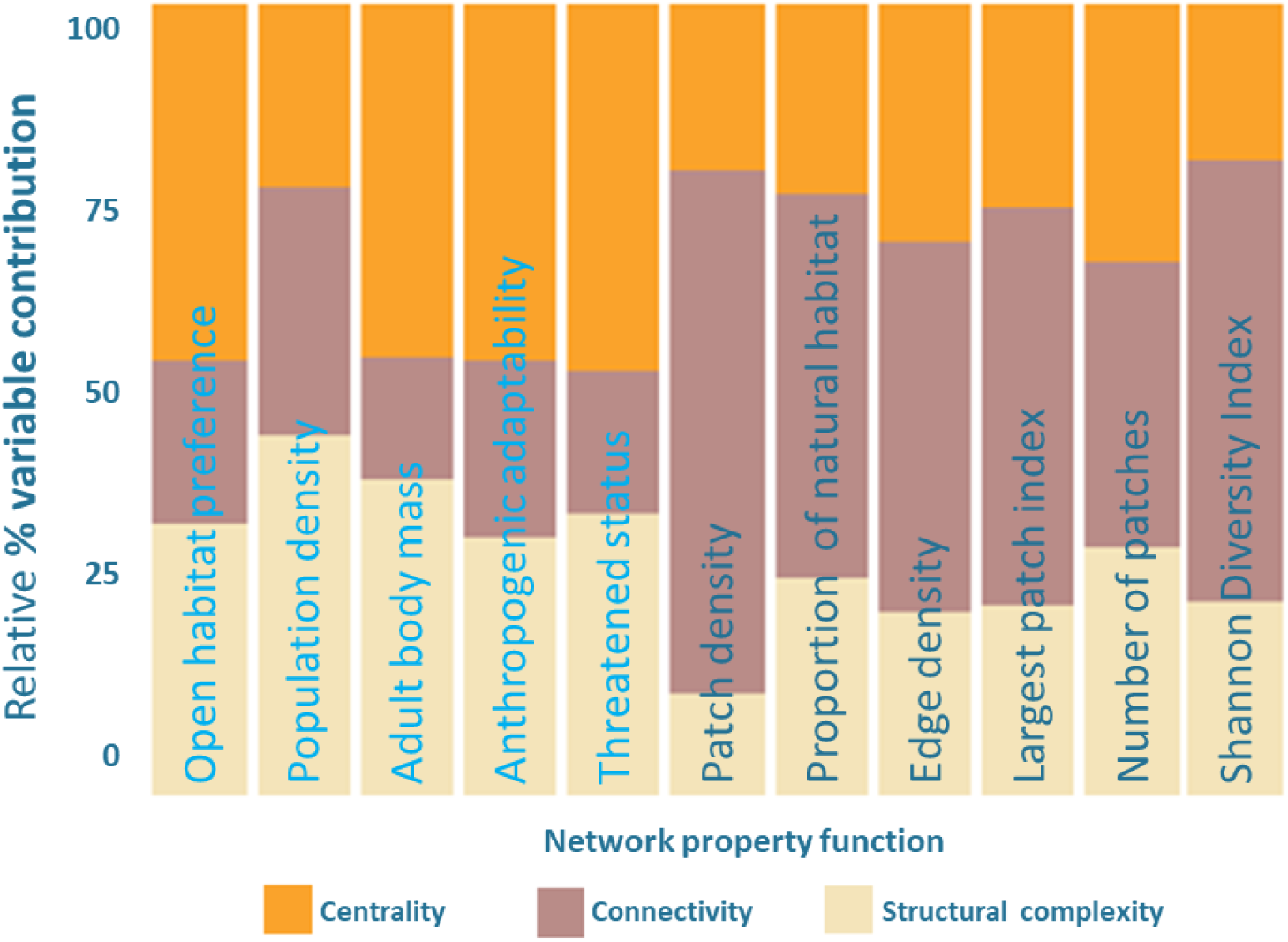
Relative variable contribution of species (light blue) and landscape (dark blue) traits in explaining network properties with different functions. Higher proportion of a colour in a bar indicates higher relative contribution of the particular variable in explaining network properties coded by that colour.

Finally, to assess the robustness of the findings, sensitivity analyses were conducted by varying the pruning threshold by ±25%. Most network metrics showed minimal variation, with stable mean values across thresholds, indicating consistency of the overall patterns. Metrics that varied significantly exhibited systematic and symmetric changes in opposite directions with increasing or decreasing thresholds, suggesting that these differences did not alter the interpretation of relative connectivity patterns (Fig. S3).

## Discussion

With increasing habitat fragmentation and population declines across broad ranges of taxonomic groups, understanding the drivers of population connectivity is paramount (Cushman, 2006). Despite advances in analytical approaches, a consensus on the factors shaping connectivity remains elusive. Connectivity has been defined in multiple ways (Merriam, 1984; Taylor et al., 1993), but overall is considered an emergent property of individual species’ dispersal capabilities and landscape structure. Although case-specific connectivity studies abound, general rules that bridge across species and landscapes remain elusive. Our study moves beyond the single-species paradigm to test whether consistent trait–connectivity associations exist in real-world landscapes. By combining species distribution models, least-cost paths, and graph-theoretic approaches, we translated complex movement pathways into comparable network metrics and evaluated them against theoretical predictions. Our results showed that while several predictions held true, particularly for landscape traits, species-level effects revealed some unexpected patterns. Body mass and threatened status were important species-level traits, with large and rare species forming dense centralized networks but with fewer nodes, suggesting importance of key habitat patches in maintaining landscape-scale connectivity. Together, these findings offer a structured framework that complements case-specific studies and advances a more general understanding of connectivity drivers across diverse carnivore assemblages.

### 4.1. High-quality patches are essential for large-bodied and threatened species

We framed predictions about species traits, particularly dispersal ability, rarity, and habitat preference. Studies have shown that multiple biological attributes are tied to body size, which can have a deep effect on the perception of landscape configuration (Marquet & Tape, 1998; Borthagaray et al., 2012). Dispersal distance, often linked to body size (Peters, 1986; Stevens et al., 2014), is widely regarded as an important determinant of connectivity (Baguette et al., 2013; Hillaert et al., 2018). As expected, larger-bodied species, although restricted to fewer habitable nodes, formed denser networks where a higher proportion of potential links were realized compared to smaller species (Gérard et al., 2021). However, these networks were also more centralized than predicted, with dependence on few high-quality nodes suggesting general vulnerability of the network.

For rare and threatened species, we expected fragmented and fragile networks (Schaffer-Smith et al., 2016), yet their networks instead resembled those of larger-bodied species - dense but centralized. This counterintuitive pattern may stem from correlations in our assemblage, where body size was positively associated with threatened status (r ∼ 0.7) and negatively with population density (r ∼ -0.6), limiting our ability to isolate the effects of rarity, like in the case of the tiger. Conversely, smaller-bodied species like the golden jackal and leopard cat are also common and least threatened. Our predictions regarding habitat preference and adaptability also received partial support. Given that over 70% of Indian land is dominated by agriculture, structurally similar to open habitats such as grasslands and scrublands, we expected open-habitat species to form resilient and redundant networks by exploiting modified landscapes. We found that the species adapted to open habitats occupy extensive, fragmented, and modular networks with many loosely connected patches, showing flexibility but low spatial cohesion. Hence, while these species did show redundant networks, they were sparse and maintained primarily by weak connections, highlighting a different form of vulnerability than anticipated.

### 4.2. Fragmentation universally lowers network efficiency

Our landscape trait predictions revolved around testing the association of contiguous natural habitats, fragmented landscapes, and heterogeneous landscapes with connectivity patterns. Unlike species effects, landscape traits showed remarkable consistency, reinforcing theoretical predictions (as outlined in Table 3). As expected, contiguous natural landscapes supported highly efficient networks with fewer isolated nodes. In addition to our prediction, we found that such networks are maintained by large, high-quality patches. There exists multiple evidence, theoretical and empirical, suggesting fragmentation and conversion of natural habitats leading to reduced species connectivity across taxa (Goodwin & Fahrig, 2002; Macdonald et al., 2018). Number of patches, edge density, and patch density have been identified as strong metrics for fragmentation and were associated with species connectivity more strongly than most other metrics tested (Cushman et al., 2013; Wasserman et al., 2013; Chambers et al., 2016). In line with both our predictions and broader empirical evidence, we found that the fragmented landscapes showed modular structures with many isolated subnetworks and reduced efficiency (r² ∼ 0.5), regardless of species identity. Finally, heterogeneous landscapes also behaved as predicted, supporting robust but spatially complex networks, underscoring the importance of matrix composition in shaping connectivity outcomes.

### 4.3. Species traits drive network structure while landscape traits determine connectivity

We found a clear division in the explanatory power of species determining structure, whereas landscape traits influenced function. Species traits contributed most to node-level properties and metrics describing network structure and centrality. In particular, open-habitat preference had a very high relative contribution to node-level properties. However, species traits contributed very little to properties that capture overall connectivity, such as network efficiency or diameter. These findings were supported by other studies that highlighted strong species effects at local scales (Borthagaray et al., 2012). Interestingly, although multiple studies suggest body size as the strongest determinant of how animals perceive the landscape (Hillaert et al., 2018; Rodríguez-Tricot & Arim, 2020; Gérard et al., 2021), and hence should determine connectivity, adult body mass was not the strongest contributor in our models. However, it did meet our predictions with denser networks for larger-bodied species.

In contrast, landscape traits were the dominant drivers of properties estimated at the scale of entire networks. An exception was landscape heterogeneity, which strongly influenced node-level metrics (Figure 3). Classic indicators such as edge density, patch density and largest patch index, widely used to categorize landscapes, were also among the strongest contributors to models explaining connectivity. Similar trends have been observed in empirical studies (Cushman et al., 2012; Cushman et al., 2013; Ash et al., 2020). Together, these findings underscore the decisive role of the landscape matrix in shaping population connectivity across species with diverse traits.

### 4.4. Graph-theoretic insights for generalizable, landscape-focused conservation

Graph theory provides a powerful framework to condense the complexity of population connectivity into simple spatial graphs with quantitative and comparable properties (Urban & Keitt, 2001). In landscape ecology, this framework has been widely adopted to evaluate network structure and the influence of landscape configuration on connectivity (Borthagaray et al., 2012; Hashemi & Darabi, 2022). Yet, the abstraction of connectivity into networks inevitably simplifies ecological reality. Local contexts, such as variation in population density, dispersal distances, or behavioral responses, are not fully captured. Our sensitivity analysis confirmed the influence of dispersal-distance dependent pruning thresholds, consistent with earlier studies (Borthagaray et al., 2012). While intraspecific differences remain important (Kanagaraj et al., 2013), our objective here was to identify patterns that generalize across species and landscapes.

Advanced methods such as circuit theory (McRae et al., 2008), resistance kernels (Compton et al., 2007), and factorial least-cost paths (Cushman et al., 2009) provide more nuanced and realistic case-specific estimates of connectivity and have been widely used for applied conservation (Atzeni et al., 2024; Lumia et al., 2024). Our intention is not to replace such approaches, but to complement them by offering a broader theoretical understanding of how species and landscapes jointly shape connectivity. The relatively modest explanatory power of our models underscores the importance of species- and context-specific differences, which should remain central to local conservation planning. Nevertheless, by moving beyond single-species and site-specific studies, our work contributes to filling a key gap in the literature - providing a framework for identifying generalizable patterns of connectivity across carnivore assemblages and landscapes.

Our findings highlight a critical conservation message - while species traits influence the structure of ecological networks, it is the configuration of landscapes that ultimately governs population connectivity. This distinction matters because conservation strategies have historically emphasized traits such as dispersal ability or habitat preference, often overlooking the overriding role of landscape composition and configuration. By disentangling the contributions of species and landscape traits, we demonstrate that maintaining large, high-quality habitat patches and managing the intervening matrix are essential levers for safeguarding connectivity across taxa.

The recent Convention on Biological Diversity (CBD) targets highlight the need for maintaining or enhancing ecological connectivity. This call for action requires prompt and practical steps to restore biodiversity by restoring and maintaining connectivity. Insights from our study underscore the need to reframe connectivity conservation from a narrow, species-centric lens to one that explicitly incorporates landscape context. Such a perspective not only strengthens our ability to design robust and generalizable conservation strategies, but also ensures that efforts remain effective under changing ecological and land-use scenarios. Emphasizing the role of landscapes in shaping connectivity, our results provide both theoretical advancement for sustaining population persistence in fragmented and modified environments.

## Supporting information

Summplementary Material

## Acknowledgement

DR is supported by the TIFR-NCBS graduate program. Our gratitude extends to the National Centre for Biological Sciences for their institutional support as facilitated by UR. The NCBS data cluster, supported under project no. 12-R&D-TFR-5.04-0900 by the Department of Atomic Energy, Government of India, was indispensable. We would like to thank Dr. Prachi Thatte and Dr. Eduardo Eizirik for their critical inputs while conceptualizing the study. Last but not the least, DR would like to acknowledge the help of multiple students who assisted in collation of data from published literature, special mention to Garima Borah, Rahul Manshani, Nikshep Trinetra, and Sunny Nama who led this gigantic task.

## Author Contributions

DR led project conceptualization, data collation, analyses, and manuscript writing. SAC assisted in revising the analytical framework and the manuscript. UR assisted in conceptualization while providing critical inputs and logistical support for the project. All authors critically contributed to the drafts and approved the final version for publication. All authors were engaged in the study from the beginning to provide a broad and holistic perspective to the designing and execution of the study. The authors declare no conflicting interests.

## Notes

https://github.com/divyashreerana/publications/tree/main/Rana-et-al_2025_Landscape_matters

### Competing Interest Statement

The authors have declared no competing interest.

## References

Aiello-Lammens, M. E., Boria, R. A., Radosavljevic, A., Vilela, B., & Anderson, R. P. (2015). spThin: an R package for spatial thinning of species occurrence records for use in ecological niche models. Ecography, 38(5), 541–545.

Ash, E., Cushman, S. A., Macdonald, D. W., Redford, T., & Kaszta, Ż. (2020). How important are resistance, dispersal ability, population density and mortality in temporally dynamic simulations of population connectivity? A case study of tigers in southeast Asia. Land, 9(11), 415.

Atzeni, L., Cushman, S. A., & Macdonald, D. W. (2024). Simulation modelling demonstrates differential performance of connectivity methods in their ability to predict genetic diversity in complex landscapes. Ecological Modelling, 498, 110886.

Baguette, M., Blanchet, S., Legrand, D., Stevens, V. M., & Turlure, C. (2013). Individual dispersal, landscape connectivity and ecological networks. Biological reviews, 88(2), 310–326.

Bender, D. J., & Fahrig, L. (2005). Matrix structure obscures the relationship between interpatch movement and patch size and isolation. Ecology, 86(4), 1023–1033.

Borthagaray, A. I., Arim, M., & Marquet, P. A. (2012). Connecting landscape structure and patterns in body size distributions. Oikos, 121(5), 697–710.

Chambers, C. L., Cushman, S. A., Medina-Fitoria, A., Martínez-Fonseca, J., & Chávez-Velásquez, M. (2016). Influences of scale on bat habitat relationships in a forested landscape in Nicaragua. Landscape Ecology, 31(6), 1299–1318.

Compton, B. W., McGarigal, K., Cushman, S. A., & Gamble, L. R. (2007). A resistant-kernel model of connectivity for amphibians that breed in vernal pools. Conservation Biology, 21(3), 788–799.

Correa Ayram, C. A., Mendoza, M. E., Etter, A., & Salicrup, D. R. P. (2016). Habitat connectivity in biodiversity conservation: A review of recent studies and applications. Progress in Physical Geography, 40(1), 7–37.

Crooks, K. R., & Sanjayan, M. (Eds.). (2006). Connectivity conservation (Vol. 14). Cambridge University Press.

Csardi, G., & Nepusz, T. (2006). The igraph software. Complex syst, 1695, 1–9.

Cushman, S. A. (2006). Effects of habitat loss and fragmentation on amphibians: a review and prospectus. Biological conservation, 128(2), 231–240.

Cushman, S. A., McKelvey, K. S., & Schwartz, M. K. (2009). Use of empirically derived source-destination models to map regional conservation corridors. Conservation Biology, 23(2), 368–376.

Cushman, S. A., Shirk, A., & Landguth, E. L. (2012). Separating the effects of habitat area, fragmentation and matrix resistance on genetic differentiation in complex landscapes. Landscape ecology, 27(3), 369–380.

Cushman, S. A., Shirk, A. J., & Landguth, E. L. (2013). Landscape genetics and limiting factors. Conservation Genetics, 14(2), 263–274.

Dinerstein, E., Loucks, C., Wikramanayake, E., Ginsberg, J., Sanderson, E., Seidensticker, J., … & Songer, M. (2007). The fate of wild tigers. BioScience, 57(6), 508–514.

Fu, Wei, et al. “Characterizing the “fragmentation–barrier” effect of road networks on landscape connectivity: A case study in Xishuangbanna, Southwest China.” Landscape and urban planning 95.3 (2010): 122–129.

Gérard, M., Marshall, L., Martinet, B., & Michez, D. (2021). Impact of landscape fragmentation and climate change on body size variation of bumblebees during the last century. Ecography, 44(2), 255–264.

Goodwin, B. J., & Fahrig, L. (2002). How does landscape structure influence landscape connectivity?. Oikos, 99(3), 552–570.

Grömping, U. (2007). Relative importance for linear regression in R: the package relaimpo. Journal of statistical software, 17, 1–27.

Hallgren, W., Santana, F., Low-Choy, S., Zhao, Y., & Mackey, B. (2019). Species distribution models can be highly sensitive to algorithm configuration. Ecological Modelling, 408, 108719.

Hashemi, R., & Darabi, H. (2022). The review of ecological network indicators in graph theory context: 2014–2021. International Journal of Environmental Research, 16(2), 24.

Hesselbarth, M. H., Sciaini, M., With, K. A., Wiegand, K., & Nowosad, J. (2019). landscapemetrics: an open-source R tool to calculate landscape metrics. Ecography, 42(10), 1648–1657.

Hillaert, J., Hovestadt, T., Vandegehuchte, M. L., & Bonte, D. (2018). Size-dependent movement explains why bigger is better in fragmented landscapes. Ecology and Evolution, 8(22), 10754–10767.

Keeley, A. T., Beier, P., & Gagnon, J. W. (2016). Estimating landscape resistance from habitat suitability: effects of data source and nonlinearities. Landscape Ecology, 31, 2151–2162.

Keeley, A. T., Beier, P., & Jenness, J. S. (2021). Connectivity metrics for conservation planning and monitoring. Biological Conservation, 255, 109008.

Kanagaraj, R., Wiegand, T., Kramer-Schadt, S., & Goyal, S. P. (2013). Using individual-based movement models to assess inter-patch connectivity for large carnivores in fragmented landscapes. Biological conservation, 167, 298–309.

Koenig, S. J., & Bender, D. J. (2018). Generalizing matrix structure affects the identification of least-cost paths and patch connectivity. Theoretical Ecology, 11(1), 95–109.

Kramer-Schadt, S., Niedballa, J., Pilgrim, J. D., Schröder, B., Lindenborn, J., Reinfelder, V., … Wilting, A. (2013). The importance of correcting for sampling bias in MaxEnt species distribution models. Diversity and distributions, 19(11), 1366–1379.

Kuhn, M., & Johnson, K. (2013). Applied predictive modeling (Vol. 26, p. 13). New York: Springer.

Liczner, A. R., Pither, R., Bennett, J. R., Bowman, J., Hall, K. R., Fletcher Jr, R. J., … & Pither, J. (2024). Advances and challenges in ecological connectivity science. Ecology and Evolution, 14(9), e70231.

Liu, C., White, M., & Newell, G. (2013). Selecting thresholds for the prediction of species occurrence with presence-only data. Journal of biogeography, 40(4), 778–789.

Lüdecke, D., Ben-Shachar, M. S., Patil, I., Waggoner, P., & Makowski, D. (2021). performance: An R package for assessment, comparison and testing of statistical models. Journal of open source software, 6(60).

Lumia, G., Modica, G., & Cushman, S. (2024). Using simulation modeling to demonstrate the performance of graph theory metrics and connectivity algorithms. Journal of Environmental Management, 352, 120073.

Macdonald, E. A., Cushman, S. A., Landguth, E. L., Hearn, A. J., Malhi, Y., & Macdonald, D. W. (2018). Simulating impacts of rapid forest loss on population size, connectivity and genetic diversity of Sunda clouded leopards (Neofelis diardi) in Borneo. PloS one, 13(9), e0196974.

Marquet, P. A., & Taper, M. L. (1998). On size and area: patterns of mammalian body size extremes across landmasses. Evolutionary Ecology, 12(2), 127–139.

McRae, B. H., Dickson, B. G., Keitt, T. H., & Shah, V. B. (2008). Using circuit theory to model connectivity in ecology, evolution, and conservation. Ecology, 89(10), 2712–2724.

Merriam, G. R. A. Y. (1984). Connectivity: a fundamental ecological characteristic of landscape pattern. In Methodology in landscape ecological research and planning: proceedings, 1st seminar, International Association of Landscape Ecology, Roskilde, Denmark, Oct 15-19, 1984/eds. J. Brandt, P. Agger.

Muscarella, R., Galante, P. J., Soley-Guardia, M., Boria, R. A., Kass, J. M., Uriarte, M., & Anderson, R. P. (2014). ENM eval: An R package for conducting spatially independent evaluations and estimating optimal model complexity for Maxent ecological niche models. Methods in ecology and evolution, 5(11), 1198–1205.

Naimi, B., & Araújo, M. B. (2016). sdm: a reproducible and extensible R platform for species distribution modelling. Ecography, 39(4), 368–375.

Peters, R. H. (1986). The ecological implications of body size (Vol. 2). Cambridge university press.

Poli, C., Hightower, J., & Fletcher Jr, R. J. (2020). Validating network connectivity with observed movement in experimental landscapes undergoing habitat destruction. Journal of Applied Ecology, 57(7), 1426–1437.

Rana, D., Sartor, C. C., Chiaverini, L., Cushman, S. A., Kaszta, Ż., Ramakrishnan, U., & Macdonald, D. W. (2024). Differentially biased sampling strategies reveal the non-stationarity of species distribution models for Indian small felids. Ecological Modelling, 493, 110749.

Rodríguez-Tricot, L., & Arim, M. (2020). From Hutchinsonian ratios to spatial scaling theory: the interplay among limiting similarity, body size and landscape structure. Ecography, 43(2), 318–327.

Samad, I., Sutaria, D., & Shanker, K. (2024). Species rich but data poor: leveraging distribution modelling techniques to map cetacean occurrence in South Asian waters. bioRxiv, 2024–12.

Savary, P., Foltête, J. C., Moal, H., Vuidel, G., & Garnier, S. (2021). graph4lg: A package for constructing and analysing graphs for landscape genetics in R. Methods in Ecology and Evolution, 12(3), 539–547.

Schaffer-Smith, D., Swenson, J. J., & Boveda-Penalba, A. J. (2016). Rapid conservation assessment for endangered species using habitat connectivity models. Environmental Conservation, 43(3), 221–230.

Schippers, P., Pouwels, R., & Verboom, J. (2025). Connecting the dots: Assessing landscape connectivity algorithms for biodiversity conservation. Ecological Modelling, 507, 111185.

Soley-Guardia, M., Alvarado-Serrano, D. F., & Anderson, R. P. (2024). Top ten hazards to avoid when modeling species distributions: a didactic guide of assumptions, problems, and recommendations. Ecography, 2024(4), e06852.

Soria, C. D., Pacifici, M., Di Marco, M., Stephen, S. M., & Rondinini, C. (2021). COMBINE: a coalesced mammal database of intrinsic and extrinsic traits.

Stevens, V. M., Whitmee, S., Le Galliard, J. F., Clobert, J., Böhning-Gaese, K., Bonte, D., … & Baguette, M. (2014). A comparative analysis of dispersal syndromes in terrestrial and semi-terrestrial animals. Ecology letters, 17(8), 1039–1052.

Thatte, P., Chandramouli, A., Tyagi, A., Patel, K., Baro, P., Chhattani, H., & Ramakrishnan, U. (2019). Human footprint differentially impacts genetic connectivity of four wide-ranging mammals in a fragmented landscape. Diversity and Distributions.

Taylor, P. D., Fahrig, L., Henein, K., & Merriam, G. (1993). Connectivity is a vital element of landscape structure. Oikos, 571–573.

Tucker, M. A., Böhning-Gaese, K., Fagan, W. F., Fryxell, J. M., Van Moorter, B., Alberts, S. C., … & Mueller, T. (2018). Moving in the Anthropocene: Global reductions in terrestrial mammalian movements. Science, 359(6374), 466–469.

Urban, D. L., Minor, E. S., Treml, E. A., & Schick, R. S. (2009). Graph models of habitat mosaics. Ecology Letters, 12(3), 260–273.

Urban, D., & Keitt, T. (2001). Landscape connectivity: a graph-theoretic perspective. Ecology, 82(5), 1205–1218.

Valavi, R., Guillera-Arroita, G., Lahoz-Monfort, J. J., & Elith, J. (2022). Predictive performance of presence-only species distribution models: a benchmark study with reproducible code. Ecological monographs, 92(1), e01486.

Wade, A. A., McKelvey, K. S., & Schwartz, M. K. (2015). Resistance-surface-based wildlife conservation connectivity modeling: Summary of efforts in the United States and guide for practitioners. Gen. Tech. Rep. RMRS-GTR-333. Fort Collins, CO: US Department of Agriculture, Forest Service, Rocky Mountain Research Station. 93 p. 333.

Wang, Z., Yang, Z., Shi, H., & Han, L. (2021). Effect of forest connectivity on the dispersal of species: A case study in the Bogda World Natural Heritage Site, Xinjiang, China. Ecological Indicators, 125, 107576.

Wasserman, T. N., Cushman, S. A., Littell, J. S., Shirk, A. J., & Landguth, E. L. (2013). Population connectivity and genetic diversity of American marten (Martes americana) in the United States northern Rocky Mountains in a climate change context. Conservation Genetics, 14(2), 529–541.

Yates, K. L., Bouchet, P. J., Caley, M. J., Mengersen, K., Randin, C. F., Parnell, S., … & Sequeira, A. M. (2018). Outstanding challenges in the transferability of ecological models. Trends in ecology & evolution, 33(10), 790–802.

Zanaga, D., Van De Kerchove, R., De Keersmaecker, W., Souverijns, N., Brockmann, C., Quast, R., … & Arino, O. (2021). ESA WorldCover 10 m 2020 v100. 2021.

Zeileis, A., Cribari-Neto, F., Gruen, B., Kosmidis, I., Simas, A. B., Rocha, A. V., & Zeileis, M. A. (2016). Package ‘betareg’. R package, 3(2), 51.

