## Supplementary material for "Landscape matters: Deriving a generalizable understanding of population connectivity using empirical data and graph theory": Summplementary Material

Codes and processed data are available in DR’s public github [repository](https://github.com/divyashreerana/publications/tree/main/Rana-et-al_2025_Landscape_matters).

[Table S1](https://docs.google.com/spreadsheets/d/1Tw9P1dnWkrzikod5rzFXWOpNlhOsuWY5kAsDYLeL2Tw/edit?usp=sharing). Species records and traits

[Table S2](https://docs.google.com/spreadsheets/d/1Tw9P1dnWkrzikod5rzFXWOpNlhOsuWY5kAsDYLeL2Tw/edit?usp=sharing). Predictor variables for building SDMs

[Table S3](https://docs.google.com/spreadsheets/u/0/d/1Tw9P1dnWkrzikod5rzFXWOpNlhOsuWY5kAsDYLeL2Tw/edit). Expected trends based on predictions

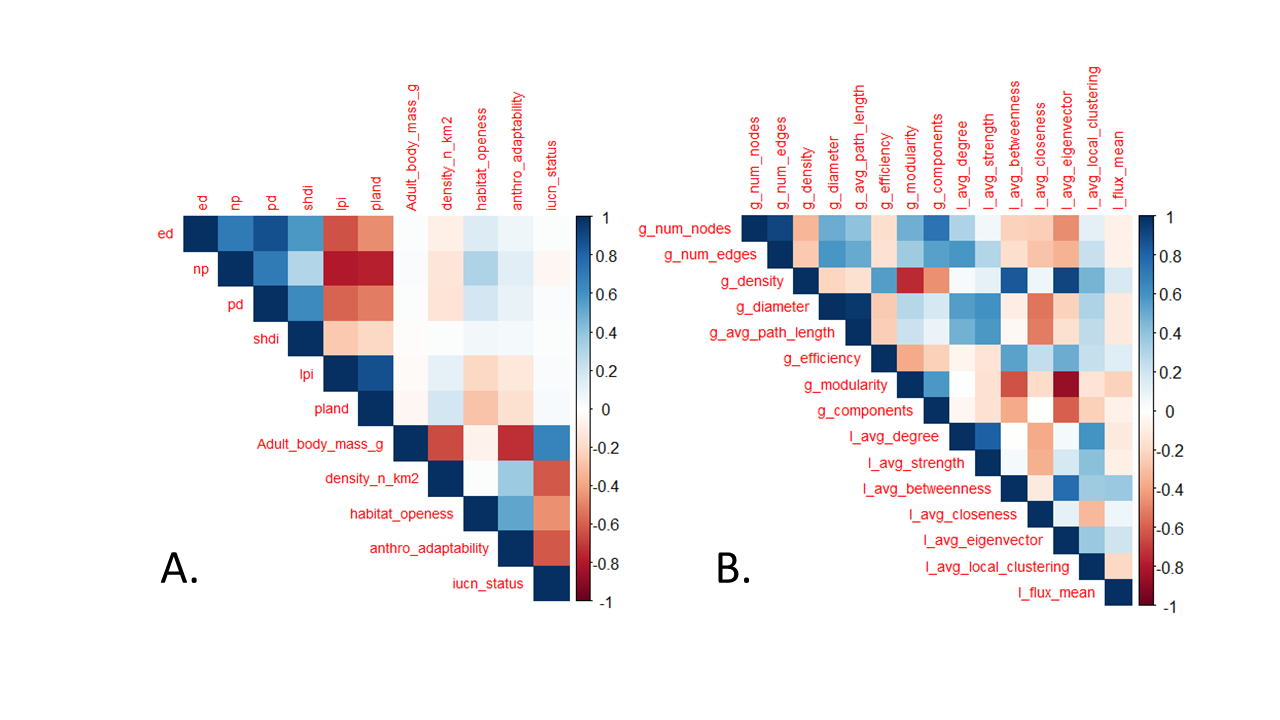

**Figure S1. Correlation plots for A. Species and landscape traits and B. Network properties**

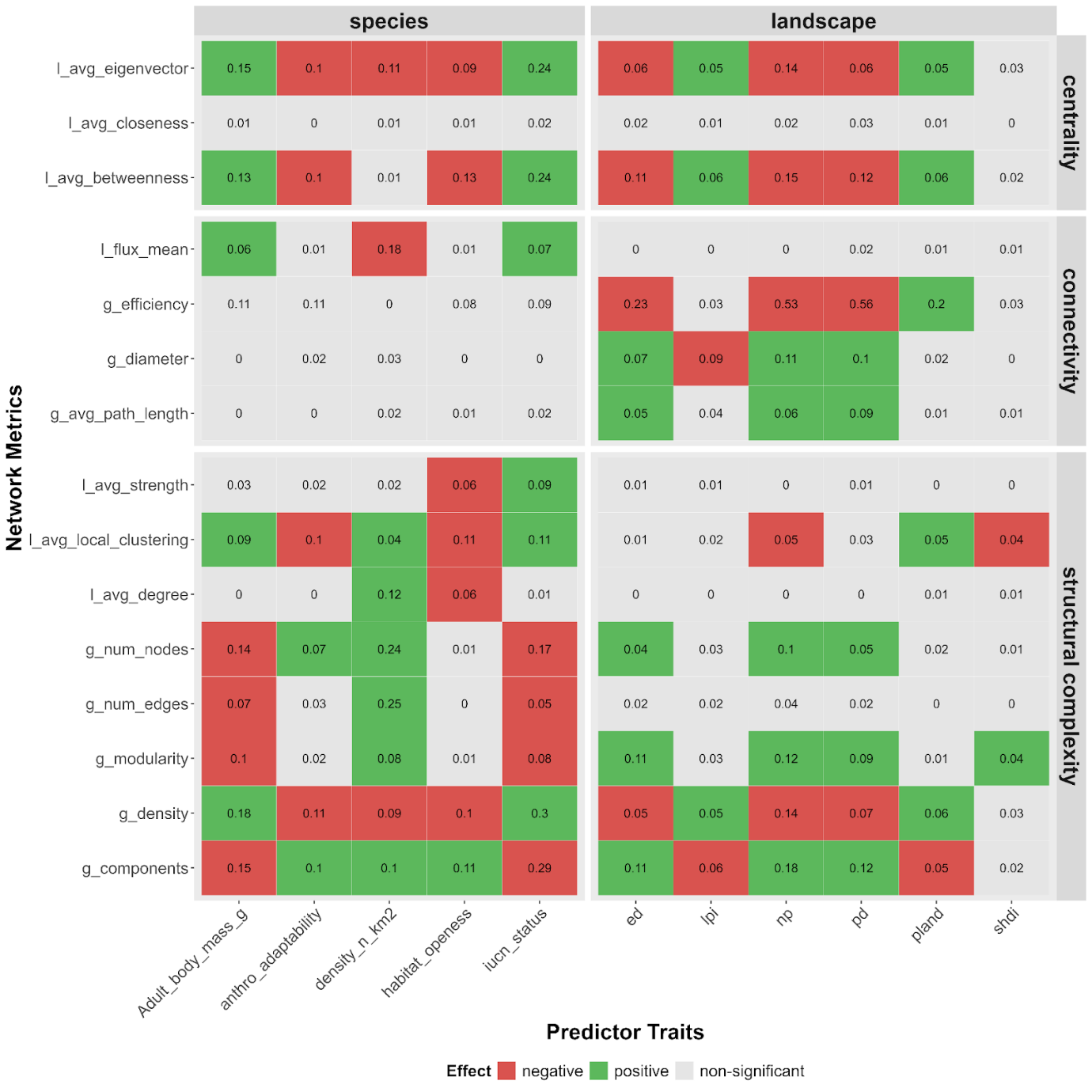

**Figure S2. Figure 2. Plot showing the results of the GLM of network properties (y-axis) with species and landscape traits (x-axis), with cells coloured for significant relations (p<0.05) in positive or negative direction. The values in each cell represent the R2 of the GLM.**

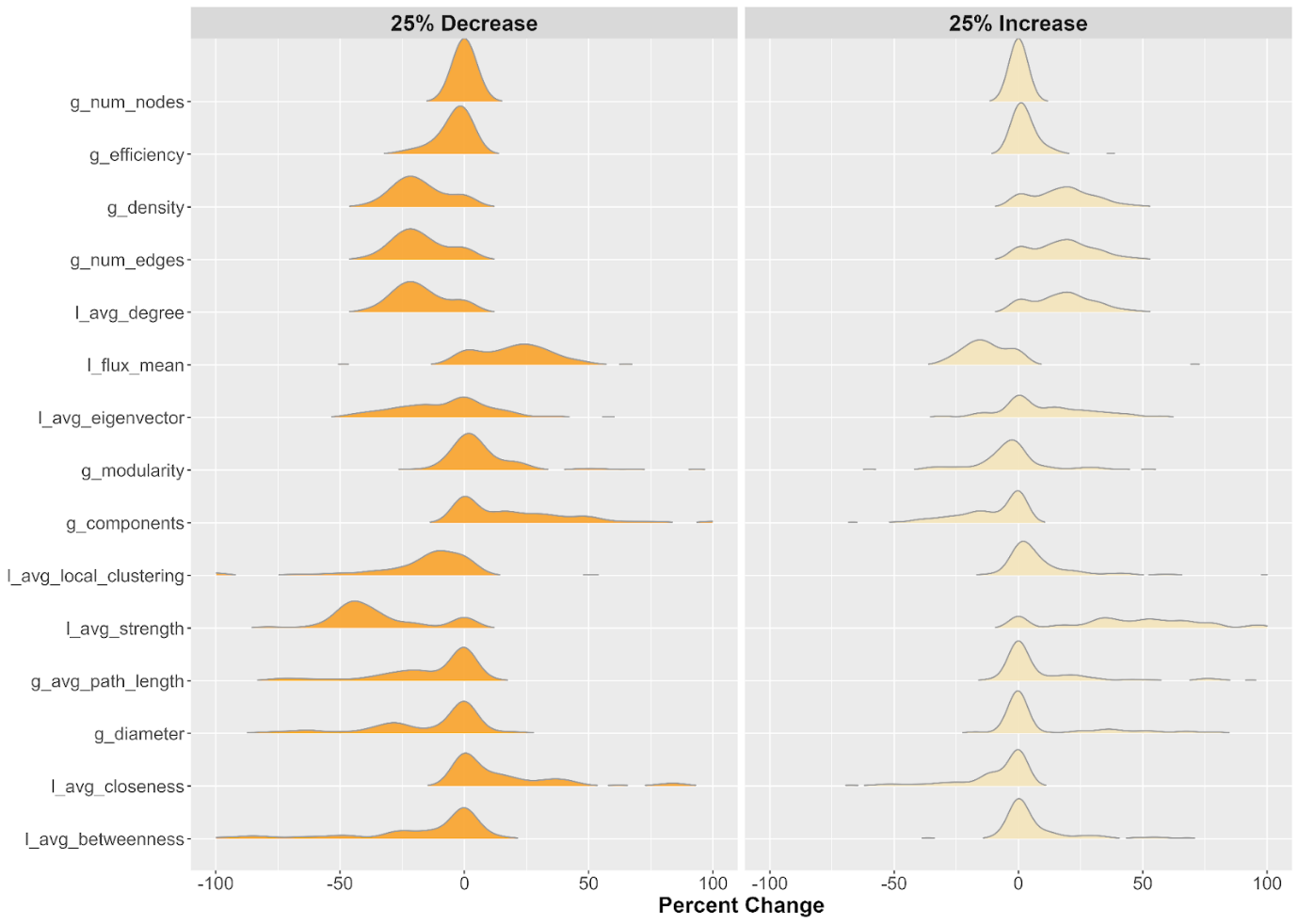

**Figure S3. Sensitivity of network properties with changes in the pruning threshold of the network. The two panels show percent change in the estimates with varying thresholds of 25% decrease and increase to the original threshold (average dispersal distance of the species). The metrics are arranged in increasing order of absolute change in the standard deviation of the estimates.**
